# Quantifying and Characterizing the Fiber in Hass Avocados During the Ripening Process

**DOI:** 10.64898/2026.04.05.716578

**Authors:** María G. Sanabria-Véaz, George C. Fahey, Knud Erik Bach Knudsen, Hannah D. Holscher

## Abstract

Reported avocado dietary fiber (DF) content and composition are inconsistently reported, particularly during ripening. Thus, this study aimed to characterize the amount and type of DF in Hass avocados and evaluate DF changes during ripening. Unripe (day 0), ripe (day 5), and overripe (day 12) Hass avocados were freeze-dried and defatted. DF was analyzed using non-starch polysaccharide (NSP) enzymatic-chemical methods. Per 100g of as-is avocado, unripe contained 3.96g total DF, ripe 3.68g, and overripe 3.26g. In ripe avocados, DF comprised 43% soluble (SDF) and 57% insoluble dietary fiber (IDF). SDF consisted primarily of rhamnogalacturonan-1 and arabinan pectins, while IDF was predominantly cellulose (32%), hemicelluloses (23%), and lignin (2%). Total DF decreased with ripening, with pectin undergoing solubilization and depolymerization, while cellulose and hemicelluloses remained stable. These findings are important as dietary fibers differentially influence intestinal microbial fermentation and health benefits.

## 1.1 INTRODUCTION

Avocados’ commercial value and consumption have increased substantially over the past 30 years (Weber & Kramer, 2022; Williams & Hanselka, 2022), which may be related to consumers’ awareness of avocado nutritional properties (Williams & Hanselka, 2022). Avocados contain many essential nutrients, including vitamins A, K, E, and C, potassium, carotenoids, and monounsaturated fatty acids (MUFA) like oleic acid (Dreher et al., 2021; Dreher & Davenport, 2013; Ford et al., 2023; Pedreschi et al., 2019). Importantly, avocados are also a rich source of dietary fiber (DF) (Dreher et al., 2021; Dreher & Davenport, 2013; Ford et al., 2023; Pedreschi et al., 2019), a nutrient that is underconsumed by ∼90% of the U.S. population (USDA & HHS, 2020). However, there are inconsistencies in the amount of DF reported in avocados, and limited information is available on the types of fibers (Pedreschi et al., 2019).

The Dietary Reference Intakes (DRIs) from the Food and Nutrition Board recommend an intake of 14g of fiber per 1000kcal/day, which corresponds to 25-38g/day for females and males, respectively. This recommendation is based on dietary fiber benefits for cardiovascular, metabolic, and digestive health (Dahl & Stewart, 2015). The United States Department of Agriculture (USDA) Food database reports 6.8g of DF per 100g of avocados (USDA, 2019), exceeding the range of 3.9g-5.5g reported by others (Li et al., 2002; Marlett, 1992; Marlett & Cheung, 1997; Wang et al., 2015). DF is mainly found in the plant cell wall and consists of pectin, cellulose, hemicelluloses, and lignin (Gill et al., 2021)

At present, the information about the type of DF in avocados is limited. There is general agreement that avocados are a significant source of pectin (Dreher & Davenport, 2013; Pedreschi et al., 2019). However, studies have focused on describing changes in pectin concentration (Defilippi et al., 2018; Huber & O’Donoghue, 1993), pectin esterification, depolymerization, and solubilization (Huber & O’Donoghue, 1993; Sakurai & Nevins, 1997), as well as enzyme activity related to pectin degradation (Awad & Young, 1979; Defilippi et al., 2018; Veau et al., 1993) during ripening, without emphasizing the type of pectin found in avocados (Pedreschi et al., 2019). Regarding hemicelluloses, there is little information on changes during ripening (Pedreschi et al., 2019). Some studies suggest xyloglucan (5-10% of edible portion) (Wakabayashi, 2000) degradation as well as little to no changes in cellulose during ripening (Sakurai & Nevins, 1997). Fruit softening related to cell wall modifications during ripening is highly variable (Brummell, 2006; Forlani et al., 2019; Wakabayashi, 2000). Ripening affects food digestibility and, thus, nutrient bioaccessibility (Holland et al., 2020). As such, this study aimed to characterize the amount and types of dietary fibers (e.g., pectin, hemicellulose, cellulose) and lignin in unripe, ripe, and overripe avocados. We hypothesized that avocado fiber composition would change during ripening.

## 1.2 METHODS

### 1.2.1 Plant Material

Hass avocados (*Persea americana*) were sourced from Michoacán, México, and supplied by West Pak, Inc. They were transported in a climate-controlled vehicle (4 °C) and received at the University of Illinois, Urbana-Champaign (UIUC) on January 31^st^, 2023. The avocados were not treated with ethylene gas and were allowed to ripen at room temperature (21 °C) and 24% relative humidity for up to 12 days. A total of 36 avocados were divided equally into three ripening stages (n=12 per group): unripe (day 0), ripe (day 5), and overripe (day 12). Ripening stages were defined based on textures and previously published criteria (Cervantes-Paz et al., 2021; Cervantes-Paz et al., 2023; Mgoma et al., 2021).

### 1.2.2 Freeze-drying

Moisture content at all ripening stages was measured using the Mettler Toledo HB43-S Series Halogen Moisture Analyzer. Avocado pulp from each ripening stage was freeze-dried using the HarvestRight Home Freeze Dryer and analyzed for moisture content to achieve < 1% residual moisture. Absolute dry matter (%DM Abs) was determined by oven-drying at 105°C. The samples were stored in a vacuum-sealed bag at −80°C until fat extraction.

### 1.2.3 Fat Extraction

Fat was extracted before fiber analysis using the Soxhlet method, following AOAC Official Method 920.39C for bulk fat extraction, as previously described (AOAC, 2002). The solvent used for the extraction was diethyl ether (500ml per 100g of dried sample). A visual summary of the ripening, freeze-drying, and fat extraction process is presented in Figure 1.

**Figure 1.**
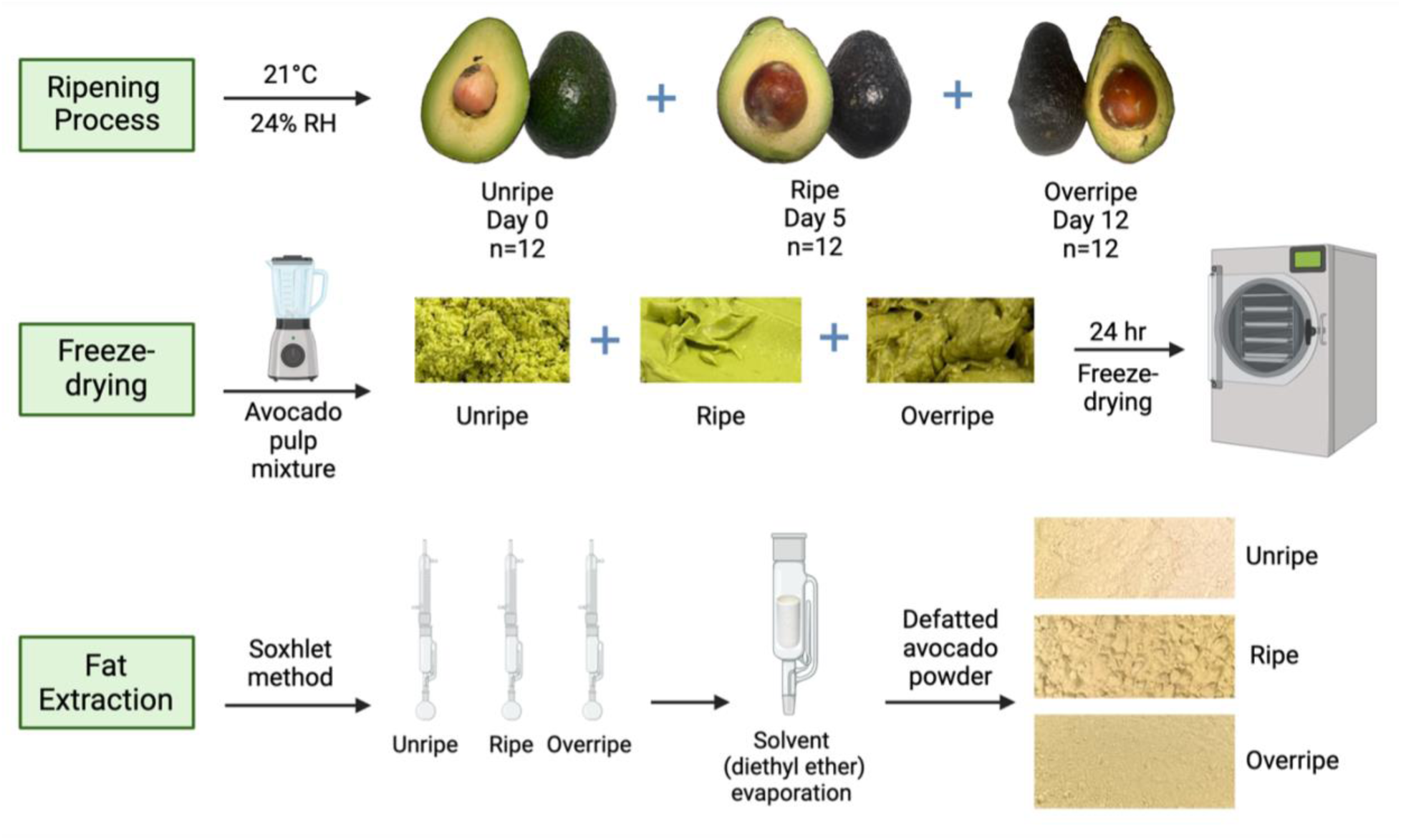
Sample processing of Hass avocados at three ripening stages for dietary fiber analysis. Avocados were ripened at room temperature (21℃) and 24% relative humidity (RH) for 12 d. The avocados from each ripening stage were mixed in a food processor and placed in a freeze-dryer for 24 h. The avocado powder (100g from each ripening stage) was placed in the Soxhlet apparatus to remove the fat.

### 1.2.4 Fiber analysis

The defatted-freeze-dried samples were shipped to Aarhus University, Denmark, for analysis of cellulose, lignin, and soluble and insoluble non-cellulosic polysaccharides (NCP) using an enzymatic-chemical-gravimetric approach (Bach Knudsen & Lærke, 2018). Modifications included omitting starch gelatinization and enzymatic degradation, and using 80% ethanol for sugar extraction. The analysis consisted of 3 procedures: A, B, and C (Figure 2). Briefly, procedures A and B measured insoluble non-starch polysaccharides (I-NSP) as neutral and acidic sugars (e.g., cellulose and total NCP residues). Cellulose content was calculated as the difference in glucose concentration between Procedure A and B. Procedure C measured the soluble NSP (S-NSP) and soluble non-cellulosic polysaccharides (S-NCP) in the sugar-free residue. Gas-liquid chromatography was used for quantification in procedures A, B, and C. Klason lignin was determined as the residue resistant to sulfuric acid hydrolysis. Total uronic acid content (galacturonic and glucuronic acid) was determined by colorimetry. All analyses were conducted in duplicate, and the reported values (refer to Supplemental Table 1 and 2) represent calculated averages.

**Figure 2.**
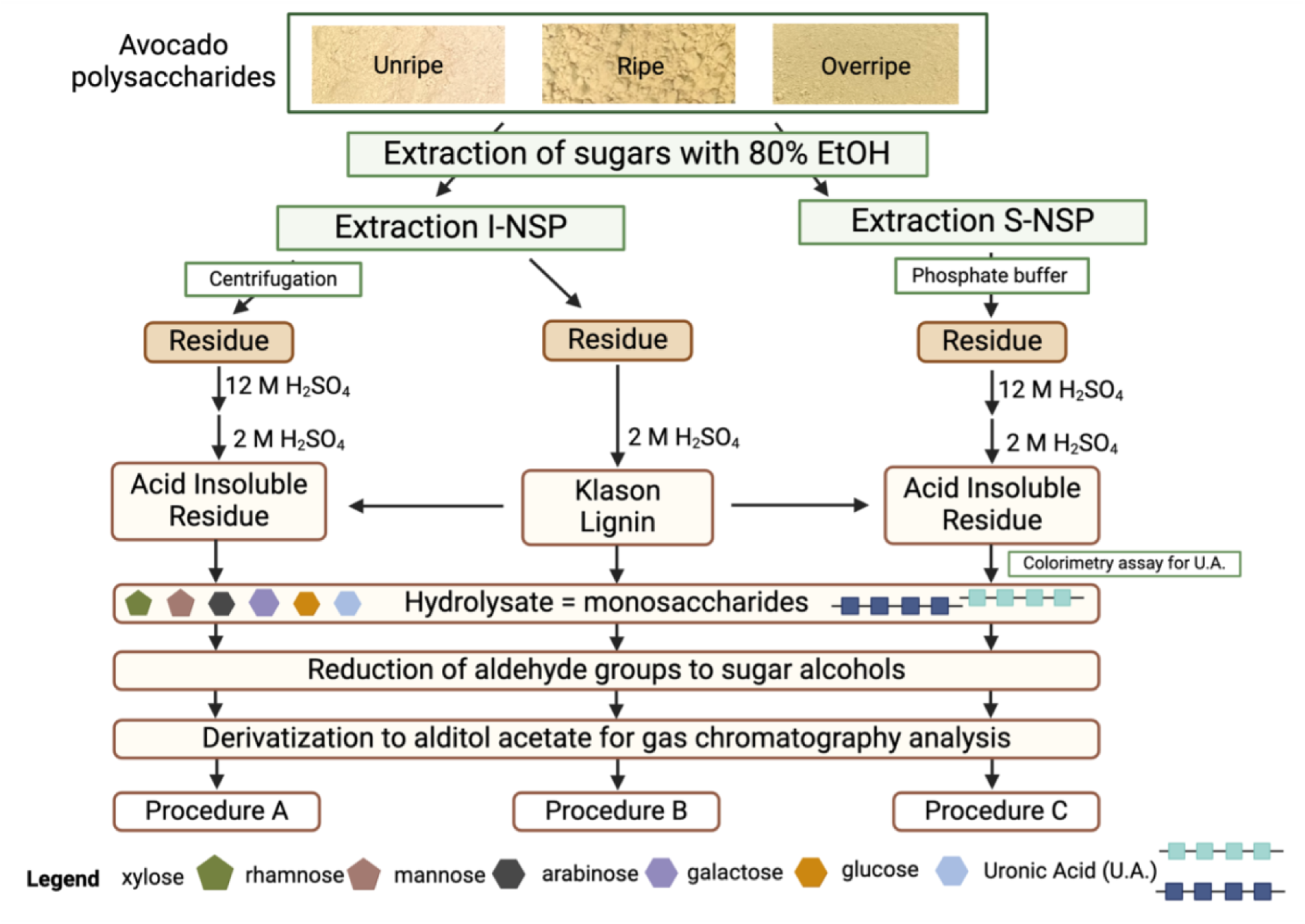
Enzymatic-chemical-gravimetric method for dietary fiber analysis. Dietary fiber was quantified using three procedures. Procedure A measures total non-starch polysaccharides (NSP) as neutral and acidic sugars (e.g., cellulose and total non-cellulosic polysaccharide residues (NCP). Procedure B measures total non-starch polysaccharides (NSP) as neutral and acidic sugars (e.g., total non-cellulosic polysaccharide residues (NCP), omitting the swelling of cellulose with 12 M H_2_SO_4_ and directly hydrolyzing the insoluble NCP (I-NCP) with 2 M H_2_SO_4_. Procedure C measures the soluble non-cellulosic polysaccharides (S-NCP) in the sugar-free residue using a phosphate buffer.

Formulas:

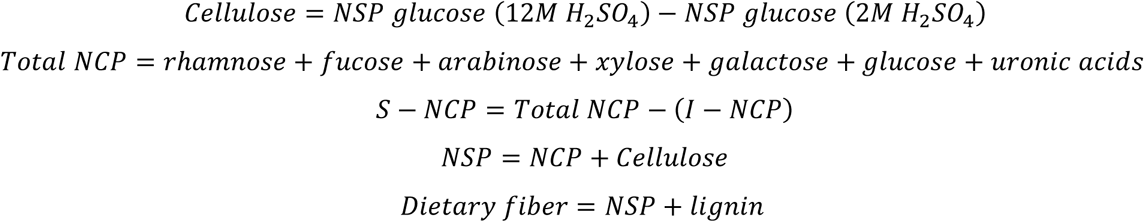

### 1.2.5 Conversion of dietary fiber values to “as-is basis”

To convert dietary fiber values from dry matter basis (DMB) to an “as-is” (wet weight) basis, moisture and fat content values (Table 1) were used. The goal was to reconstruct the composition of fresh avocado pulp by accounting for water and fat content lost during freeze-drying and fat extraction so that results would be more translatable to the composition when consumed. The samples were freeze-dried, and then a sub-sample was oven-dried (105°C) to determine dry matter content. These values were multiplied to calculate the absolute dry matter percentage (%DM Abs).

**Table 1.**
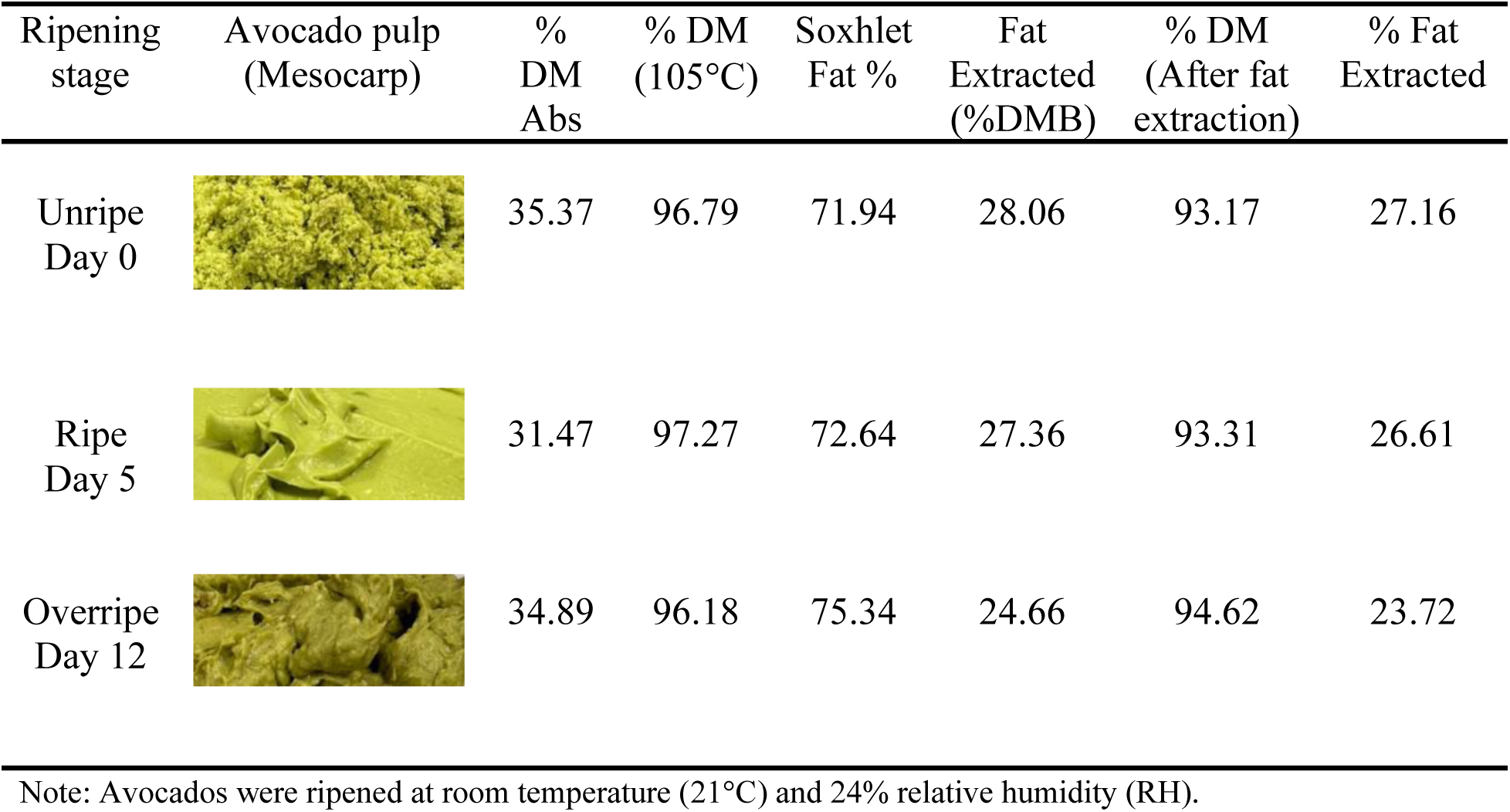
Changes in avocado pulp (mesocarp), % dry matter (%DM), and total fat at three ripening stages.

| Ripening stage | Avocado pulp (Mesocarp) | % DM Abs | % DM (105°C) | Soxhlet Fat % | Fat Extracted (%DMB) | % DM (After fat extraction) | % Fat Extracted |
| --- | --- | --- | --- | --- | --- | --- | --- |
| Unripe Day 0 |  | 35.37 | 96.79 | 71.94 | 28.06 | 93.17 | 27.16 |
| Ripe Day 5 |  | 31.47 | 97.27 | 72.64 | 27.36 | 93.31 | 26.61 |
| Overripe Day 12 |  | 34.89 | 96.18 | 75.34 | 24.66 | 94.62 | 23.72 |

Note: Avocados were ripened at room temperature (21°C) and 24% relative humidity (RH).

Then, the % Soxhlet fat was used to calculate % Fat Extracted on a DMB by taking the difference (100 − % *Soxhlet Fat*) for each ripening stage. Fat content on an “as-is” basis was calculated using the following formula: 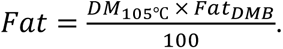

Lastly, each fiber component was converted to an “as-is” basis using the following formula:

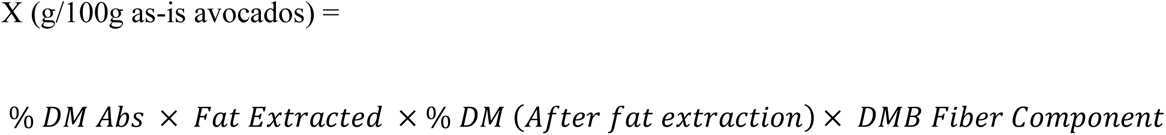

Calculations were conducted for each fiber component (e.g., NSP, NCP, cellulose, lignin, TDF) across ripening stages.

## 1.3 RESULTS AND DISCUSSION

### 1.3.1 Changes in avocado pulp (mesocarp) and moisture during ripening

Hass avocados undergo textural and color changes in the skin from green to purple/black during ripening related to changes in cell wall modifications, organoleptic properties, and chlorophyll decrease (Cox et al., 2004). The changes in green color from bright to olive green, as seen in Table 1, could be related to chlorophyll reduction and degradation, as well as enzymatic browning caused by overripening. The %DM Absolute (Abs) is an important quality standard for export, with an optimal range of 23–28% (Ford et al., 2023; Giuggioli et al., 2021). However, agricultural growing practices and varietal seasons, postharvest handling such as external ethylene treatment, oil content, and the overall ripening process affect the %DM (Ford et al., 2023; Giuggioli et al., 2021). For instance, Ozdemir and Topuz reported a steady increase in %DM of 43.6% in avocados harvested from November to January. Indeed, the Ozdemir and Topuz ripe Hass avocados harvested in January had a %DM of 31.09 ± 0.389 (Ozdemir & Topuz, 2004), which is similar to the results reported herein (31.47%; Table 1).

In agreement with our results and Ozdemir and Topuz, Méndez Hernández et al. reported %DM of 26.8%±3.48 in November and 31.4%±2.91 in February (Méndez Hernández et al., 2023). The avocados from Méndez Hernández et al. were taken from Northern areas of the Canary Islands, with a mean relative humidity (RH) and temperature of 77.5% and 19.4°C, respectively, from November to February, which is similar to the mean annual temperature in Michoacán, México, of 19.3°C (Medina-Carrillo et al., 2017). Further, %DM increases at both ambient temperatures and lower RH, which increases moisture loss and, thus, increases %DM (Kassim & Workneh, 2020). The influence of temperature during harvesting and storage at room temperature under low relative humidity likely contributed to the elevated DM values.

### 1.3.2 Dietary fiber in avocados - pectin

NCP, cellulose, NSP, lignin, and TDF data on an “as-is basis” are presented in Table 2, and NCP data on an “as-is basis” are presented in Table 3. The amount of S-NCP in ripe avocados (1.59g/100g) was the highest among the ripening stages, followed by unripe (1.40g/100g) and overripe (1.11g/100g) (Table 2). The major sugar monomers in S-NCP for the ripe avocados were uronic acid (UA) (0.96g; 60.3%), arabinose (0.39g; 24.5%), galactose (0.11g; 6.8%), and rhamnose (0.05g; 2.9%) (Table 3). This proportion of monosaccharides in ripe avocados is most likely indicative of soluble pectin.

**Table 2.**
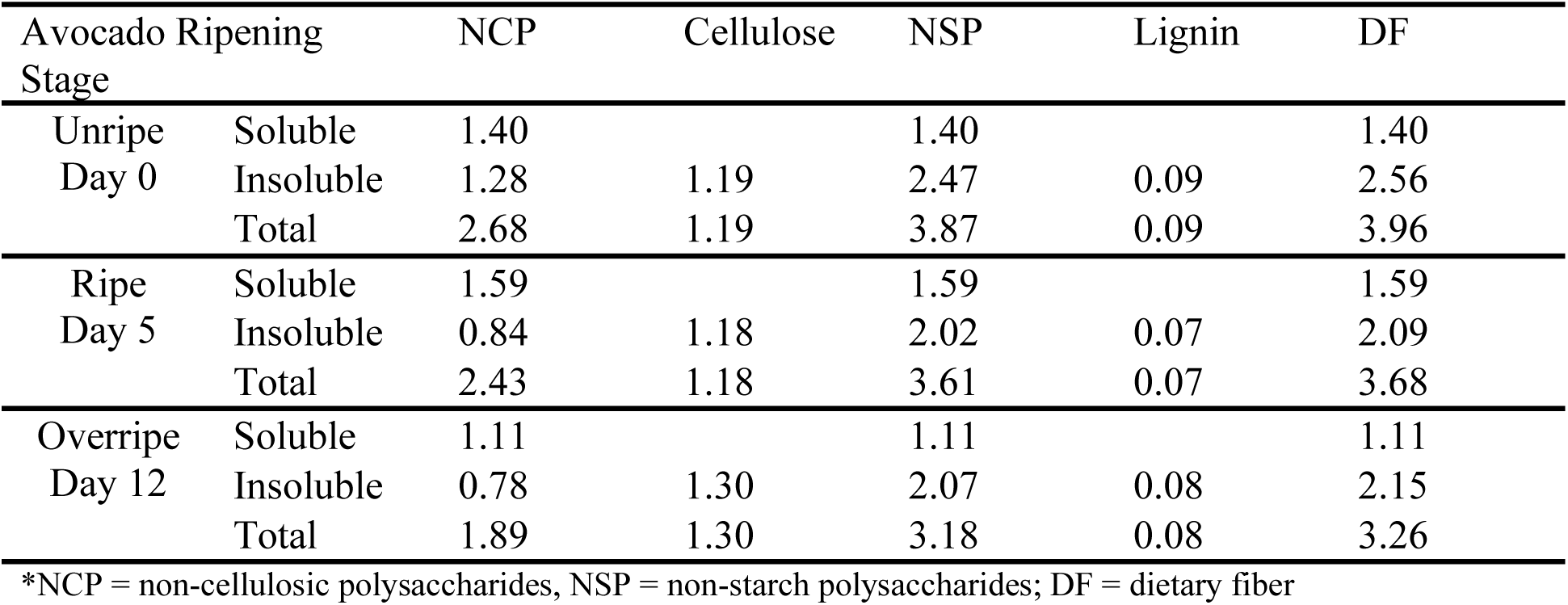
Changes in avocado fiber constituents on a wet, “as-is” basis (g/100g) at different ripening stages.

| Avocado Ripening Stage |  | NCP | Cellulose | NSP | Lignin | DF |
| --- | --- | --- | --- | --- | --- | --- |
| Unripe Day 0 | Soluble | 1.40 |  | 1.40 |  | 1.40 |
|  | Insoluble | 1.28 | 1.19 | 2.47 | 0.09 | 2.56 |
|  | Total | 2.68 | 1.19 | 3.87 | 0.09 | 3.96 |
| Ripe Day 5 | Soluble | 1.59 |  | 1.59 |  | 1.59 |
|  | Insoluble | 0.84 | 1.18 | 2.02 | 0.07 | 2.09 |
|  | Total | 2.43 | 1.18 | 3.61 | 0.07 | 3.68 |
| Overripe Day 12 | Soluble | 1.11 |  | 1.11 |  | 1.11 |
|  | Insoluble | 0.78 | 1.30 | 2.07 | 0.08 | 2.15 |
|  | Total | 1.89 | 1.30 | 3.18 | 0.08 | 3.26 |

\*NCP = non-cellulosic polysaccharides, NSP = non-starch polysaccharides; DF = dietary fiber

**Table 3.** Non-cellulosic polysaccharide (NCP) residues on a wet, as-is basis (g/100g) from soluble and insoluble fractions of avocado fiber at different ripening stages.

| Avocado Ripening Stage |  | Rha | Fuc | Ara | Xyl | Man | Gal | Glu | U.A. |
| --- | --- | --- | --- | --- | --- | --- | --- | --- | --- |
| Unripe<br>Day 0 | Soluble | 0.03 | 0.00 | 0.24 | 0.00 | 0.00 | 0.17 | 0.01 | 0.94 |
|  | Insoluble | 0.02 | 0.03 | 0.23 | 0.32 | 0.12 | 0.22 | 0.09 | 0.25 |
|  | Total | 0.04 | 0.04 | 0.47 | 0.32 | 0.12 | 0.39 | 0.10 | 1.19 |
| Ripe<br>Day 5 | Soluble | 0.05 | 0.00 | 0.39 | 0.01 | 0.01 | 0.11 | 0.08 | 0.96 |
|  | Insoluble | 0.01 | 0.03 | 0.13 | 0.32 | 0.10 | 0.11 | 0.00 | 0.15 |
|  | Total | 0.05 | 0.03 | 0.52 | 0.31 | 0.11 | 0.22 | 0.08 | 1.11 |
| Overripe<br>Day 12 | Soluble | 0.05 | 0.00 | 0.32 | 0.02 | 0.01 | 0.11 | 0.09 | 0.51 |
|  | Insoluble | 0.01 | 0.03 | 0.09 | 0.31 | 0.11 | 0.11 | 0.00 | 0.12 |
|  | Total | 0.06 | 0.03 | 0.41 | 0.32 | 0.12 | 0.22 | 0.09 | 0.63 |

\*Rha=rhamnose, Fuc=fucose, Ara=arabinose, Xyl=xylose, Man=mannose, Gal=galactose, Glu=glucose, U.A.=uronic acid

Pectins are a heterogeneous group of soluble polysaccharides containing galacturonic acid residues, varying degrees of polymerization, esterification, and sugar side chains (Prasanna et al., 2007; Shi et al., 2023). The main types of pectin are homogalacturonan (HG), which is a linear polymer of α-(1,4)-d-galacturonic acid, and rhamnogalacturonan types 1 and 2 (RG-1 and RG-2), which are mostly branched with rhamnose residues in the HG chain. Other types of pectic polysaccharides, such as arabinans, galactans, and arabinogalactans, are considered neutral pectins because they lack galacturonic acid and can be found as free polymers or as side chains of RG-1 (Prasanna et al., 2007; Spadoni Andreani et al., 2021).

The monosaccharide composition (i.e., rhamnose, arabinose, and galactose) in the S-NCP suggests the presence of rhamnogalacturonan pectin and a lack of homogalacturonan (Hachem et al., 2016). Rhamnogalacturonan-1 (RG-1) has been associated with avocado ripening (Pedreschi et al., 2019) and can be distinguished from homogalacturonan by the ratio of uronic acid to rhamnose (UA/Rha). A higher ratio indicates a higher proportion of homogalacturonan (Basanta et al., 2014), whereas a lower ratio indicates a higher proportion of RG-1. A study by Brahem et al. characterized the tissue-specific differences in cell wall polysaccharides of ripe and overripe pears by the UA/Rha ratio (Brahem et al., 2017). The researchers obtained ratios of 17 and 6 in the water-soluble fraction of ripe and overripe pears, respectively, which indicates the presence of highly retained rhamnogalacturonan (Brahem et al., 2017). Similarly, pectins from ripe olives had a higher ratio of Rha/UA than unripe olives, i.e., pectins in the ripe olive fruit are more branched (Vierhuis, 2002). In this current study, the ratio of UA/Rha in the S-NCP of ripe avocados was 20.5, compared to 35.0 in unripe and 9.4 in overripe (Figure 3), which indicates a higher abundance of rhamnogalacturonan-1 compared to homogalacturonan.

**Figure 3.**
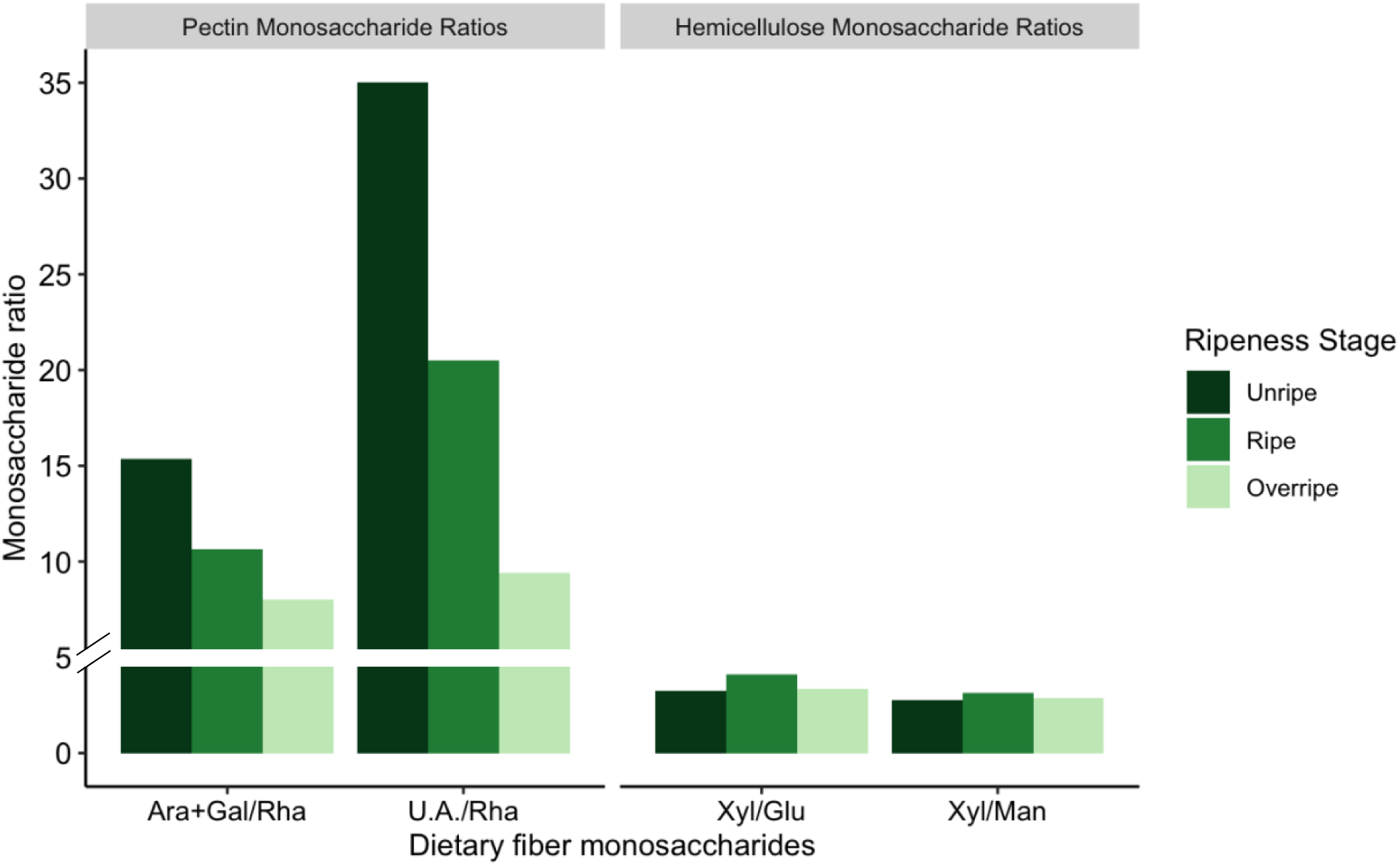
Monosaccharide ratios to analyze the degree of branching and depolymerization of pectin and hemicellulose at different degrees of ripening. Monosaccharide ratios are calculated using the concentration of uronic acid, rhamnose, arabinose, galactose, xylose, mannose, and glucose across ripening stages. Ratios of uronic acid/rhamnose (UA/Rha) and arabinose+galactose/rhamnose (Ara+Gal)/Rha are characteristic of pectin. Ratios of xylose/mannose (Xyl/Man) and xylose/glucose (Xyl/Glu) are characteristic of hemicelluloses.

Further, the abundance of arabinose relative to galactose can be indicative of side chains such as arabinans and/or arabinogalactans (Aboughe-Angone et al., 2008; Basanta et al., 2014; Hachem et al., 2016), previously found in ripe, ready-to-eat avocados (Marlett, 1992). To analyze this, we calculated the ratio of arabinose+galactose to rhamnose [(Ara+Gal/Rha)] as an indication of neutral side chains in the rhamnogalacturonan-1 backbone. A higher ratio is related to greater branching and/or differences in relative proportions of homogalacturonan to rhamnogalacturonan-1. In this report, the ratio of Ara+Gal/Rha was higher in unripe (15.3), followed by ripe (10.6) and overripe (8.0) (Figure 3). The low UA/Rha and Ara+Gal/Rha ratio further indicates rhamnogalacturonan-1 pectin (Parkar et al., 2021), with fewer side chains and a degree of branching found in ripe avocados. Also, the lower Ara+Gal/Rha ratio in overripe avocados indicates removal of rhamnogalacturonan-1 side chains during ripening (Brahem et al., 2017). Therefore, the main components of the SDF fraction are the pectin rhamnogalacturonan-1 with side chains of arabinans and/or arabinogalactans, as evidenced by the presence of uronic acids, rhamnose, arabinose, and galactose.

### 1.3.3 Pectin changes during ripening

Indications of increased solubility of uronic acid during ripening (Huber & O’Donoghue, 1993) as well as degradation to lower molecular size (Sakurai & Nevins, 1997), potentially through pectin-degrading enzymes (e.g., pectinerase and polygalacturanase), have been reported in avocado fruits. Our work indicated that the solubility of uronic acid increased from unripe (10.5%) to ripe (12.3%) avocados, representing a 17% increase (Supplemental Table 2). Similarly, a study by Sakurai and Nevins reported a 32% increase in uronic acid content of the water-soluble fraction from unripe (day 0) to fully ripe (day 6) avocados, while soluble uronic acid content in the ripe avocados represented 86.6% of the total uronic acid content (Sakurai & Nevins, 1997). In addition to increased uronic acid solubility in ripe avocados, rhamnogalacturonan-1 undergoes modifications in the side chains composed of galactose (e.g., galactans, arabinogalactans) by beta-galactosidase and the solubilization of arabinose (Brummell, 2006; Brummell & Harpster, 2001; Defilippi et al., 2018; Veau et al., 1993).

In the current study, galactose content in the S-NCP decreased by 54.5%, whereas arabinose content in the S-NCP increased by 62.5% (Table 3). The rhamnose content remained constant throughout the ripening stages (0.04-0.06g), suggesting that changes in the pectin molecule are mostly related to changes in the side chains (Pedreschi et al., 2019). Interestingly, a study by Defilippi and colleagues reported increased beta-galactosidase activity in ripe avocados, paralleled by a 50% decrease in galactose and an increase in arabinose in water-soluble pectin (Defilippi et al., 2018). Likewise, in a study by Basanta et al evaluating changes in cell wall polysaccharides in sweet cherries during ripening, there was a reduction of galactose and a concomitant increase in arabinose (Basanta et al., 2014).

### 1.3.4 Dietary fiber in avocados – hemicelluloses

The Insoluble Dietary Fiber (IDF) is the sum of the hemicelluloses (I-NCP), cellulose, and lignin (Table 2). Overall, unripe avocados contained the highest amount of I-NCP (1.28g/100g), followed by ripe (0.84g/100g), and overripe (0.78g/100g) (Table 2). The major sugar monomers in the I-NCP fraction of ripe avocados were xylose (0.32g; 38.1%), arabinose (0.13g; 15.5%), galactose (0.11g; 13.1%), mannose (0.10g; 12%), and fucose (0.03g; 0.4%) (Table 3).

Hemicelluloses are insoluble, non-pectic polysaccharides in plant cell walls, primarily composed of neutral sugars like xylose, glucose, mannose, galactose, and arabinose (An et al., 2022; Decker et al., 2008). The main types of hemicelluloses include xylans, mannans, beta-glucans, and xyloglucans (Brummell, 2006; Padayachee et al., 2017). The monosaccharide composition (i.e., xylose, glucose, galactose, fucose, and arabinose) in the I-NCP is indicative of xyloglucan (O’Donoghue & Huber, 1992; Park & Cosgrove, 2015; Sakurai & Nevins, 1997), which is the primary type of hemicellulose (∼ 20-25%), found in the primary cell wall of dicot plants like avocados. Xyloglucan consists of a 1,4-β-D-glucan backbone (G), with 1,6-α-D xylose moieties every three glucose units and potential links with galactose and/or fucose.

The major presence of xyloglucans in dicot plants generally contributes to a higher ratio of xylose to glucose, compared to monocots (Vierhuis, 2002). A ratio of xylose to glucose (Xyl/Glu) of 0.75 is suggested for xyloglucan in higher plants (Aboughe-Angone et al., 2008). A higher ratio can be due to the type of plant (dicots vs. monocots) or the presence of hemicelluloses other than xyloglucan (Basanta et al., 2014). Figure 3 in this paper shows that the ratio of Xyl/Glu in ripe avocados is 4.1. Similarly, Aboughe-Angone et al. (Aboughe-Angone et al., 2008) reported a ratio of 3.9 in the alkali-soluble fractions of argan fruit. The study concluded that the fruit contains xyloglucan and a mix of other hemicellulosic polysaccharides. Thus, we calculated the ratio of xylose to mannose (Xyl/Man) in the I-NCP, which describes the contribution of xylans or mannans. The ratio of Xyl/Man in ripe avocados was 3.15 (Figure 3), indicating a higher degree of xylans than mannans. The presence of galactose in the insoluble fraction could be related to the xyloglucan structure. However, the presence of arabinogalactans cannot be ruled out, possibly related to side-chain interactions between hemicelluloses and RG-1 structures in the cell wall (Basanta et al., 2014). In summary, the ripe avocados in this report contain xyloglucans, but other polysaccharides (e.g., xylan, arabinogalactans) may also contribute to glycosidic linkages in the hemicelluloses of avocados.

### 1.3.5 Hemicellulose changes during ripening

As observed with pectin in S-NCP, hemicelluloses are potentially subject to depolymerization and enzyme degradation (Brummell, 2006), particularly by cellulase (Decker et al., 2008; Wakabayashi et al., 2000). One of the first studies examining hemicelluloses in avocados during ripening reported depolymerization caused by cellulase to lower molecular weight hemicelluloses but with no specific relation to xyloglucan (O’Donoghue & Huber, 1992). However, other studies reported a molecular weight reduction of xyloglucan in avocados with no changes in the individual sugar components of hemicelluloses, particularly xylose, mannose, and glucose (Sakurai & Nevins, 1997).

In the present study, the amount of xylose and mannose in the I-NCP stayed relatively constant (Table 3). As seen in Figure 3, the ratios of Xyl/Glu and Xyl/Man were slightly higher in ripe avocados. However, compared to the changes observed in pectin, the ratios in the I-NCP remained relatively similar at each ripening stage, suggesting little to no changes in xylan and/or xyloglucan structure during ripening. Further, the 50% decrease in galactose (Table 3) may reflect reductions in arabinogalactans linked to rhamnogalacturonan-1 side chains (Brummell, 2006). Overall, the stable sugar ratio profile and possibly lower content of hemicelluloses from 1.28 (unripe) to 0.78g (overripe) (Table 2)], during ripening, particularly xyloglucan, suggests changes in interactions with other fiber components (i.e., cellulose, pectin) and potentially breakdown of cell wall matrix, rather than structural changes within hemicellulose.

### 1.3.6 Dietary fiber in avocados – cellulose

The highest amount of cellulose (g/100g) was seen in overripe avocados (1.30g), followed by unripe (1.19g) and ripe (1.18g) (Table 2), with cellulose representing 48%, 58%, and 63% of insoluble-NSP (I-NSP) in unripe, ripe, and overripe avocados, respectively. Cellulose is a linear, tightly packed, insoluble polysaccharide made of glucose units linked by β-(1➔4) linkages and with hydrogen bonds between adjacent chains (Padayachee et al., 2017). As part of the cell wall, cellulose disassembly may contribute to fruit softening, particularly due to increased cellulase (endoglucanase) (El-Zoghbi, 1994) activity followed by ethylene production characteristic of climacteric fruits (Kou et al., 2021; Paul et al., 2012).

Questions have arisen regarding the role of cellulase in ripening, considering that avocado mesocarp produces significant amounts of this enzyme (Awad & Young, 1979; Sakurai & Nevins, 1997). Some studies report higher cellulase activity with decreased fruit firmness (Pesis et al., 1978) and potential decrease in cellulose content (El-Zoghbi, 1994), whereas other studies do not report a correlation between cellulose content and fruit firmness (D. Li et al., 2022), which may be related to differences in fruits and cultivars. Aside from its potential role in xyloglucan and cellulose breakdown, another possible role of cellulase is its impact on the proportion of crystalline cellulose by disrupting the peripheral and amorphous (less tightly packed) cellulose microfibril regions (Brummell, 2006; O’Donoghue et al., 1994). A study by O’Donoghue et al. using transmission electron microscopy with cytochemical localization showed that cellulose changes fibril organization, rather than complete breakdown as typically seen with pectin and/or hemicelluloses (O’Donoghue et al., 1994). Further, as the fruit ripens, the cellulose content shifts from the middle lamella to the cell wall periphery. This results in the same amount of cellulose through ripening during fiber analysis.

In this study, cellulose content remained stable from 1.19g to 1.30g per 100g of fresh avocados. However, as a percentage of I-NSP, there is a slight increase in cellulose with ripening, which could be related to the disruption of crystalline cellulose to less tightly packed molecules and/or changes in hydrogen bonding with hemicelluloses (Brummell, 2006; O’Donoghue et al., 1994; Pedreschi et al., 2019). Overall, cellulose was present in high amounts in ripe avocados, and the content remained relatively stable throughout the ripening stages (Table 2).

### 1.3.7 Dietary fiber in avocados – lignin

Our work indicated that the lignin content (g/100g) remained constant at all ripening stages: unripe (0.09g), overripe (0.08g), and ripe (0.07g), representing 3.5%, 3.7%, and 3.3%, respectively, of IDF (Table 2). Lignin is a rigid polymer that provides cell structure and is related to plant tissue firmness and texture (Annunziata, 2019). The changes in lignin content during ripening, particularly in the mesocarp, are not well studied compared to other cell wall polysaccharides (Medina-Carrillo et al., 2017). As lignin is related to structural components, higher lignin concentrations are expected in unripe fruits, and degradation is expected as the fruit ripens. Further research is needed to explore the role of lignin in the ripening of avocados and other climacteric fruits.

### 1.3.8 Total Dietary Fiber (TDF)

Overall, the highest amount of TDF (g/100g) was observed in unripe avocados (3.96g), followed by ripe (3.68g) and overripe (3.26g) (Table 2), indicating a decreasing trend in fiber amounts as the fruit ripened. This has been reported in other fruits such as mangoes, guava, dates, and strawberries (El-Zoghbi, 1994) and could be related to the extensive hydrolysis caused by cell-wall degrading enzymes such as pectinesterase, β-galactosidase, polygalacturonase, and cellulase, which have been documented in avocados and discussed previously. Further, ripe avocados contain a mixture of 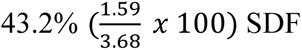 and 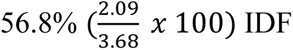 (Table 2).

Our results are in agreement with Lund et al., 1983—the amount of cellulose, SDF, and IDF in ripe avocados obtained by these researchers (1.16g±0.007, 1.29g±0.07, 2.04g±0.03, respectively) was similar to the results reported herein (1.18g, 1.59g, 2.09g, respectively, Table 2). However, the insoluble hemicellulose and lignin (1.32g±0.10 and 0.18g±0.016, respectively) were slightly higher than in the present study (0.84g and 0.07g, respectively). These differences could be related to the avocado variety (tropical vs. subtropical) and fat extraction method (e.g., length and solvent used (Ford et al., 2023). In another study by (Marlett, 1992), ripe avocados contained 1.3g/100g of SDF and 2.6g/100g of IDF. Overall, the SDF reported by Marlett et al. was similar in our study and in the work of (Lund et al., 1983), whereas IDF was slightly higher. Nonetheless, of the 2.6g of IDF reported by (Marlett, 1992), 1.4g were exclusively cellulose, which is very similar to the values obtained by Lund et al. and the results reported herein. Lastly, a study by Sánchez-Castillo et al., examining NSP in Mexican fruits and vegetables, compared the DF from Hass and “criollo” avocado cultivars, among other foods (Sánchez-Castillo et al., 1995). The researchers employed a similar method to that in the present study, revealing that the TDF was comparable between (Sánchez-Castillo et al., 1995) and our work (3.6g vs. 3.68g, respectively). Furthermore, Sánchez-Castillo et al. reported values of 1.6g/100g of SDF and 2g/100g of IDF, similar to the values in our work (1.59g SDF and 2.09g IDF; Table 2).

Notably, some studies have reported higher DF values than those observed in the present study and others (Lund et al., 1983; Marlett, 1992; Marlett & Cheung, 1997; Sánchez-Castillo et al., 1995). For instance, Wang et al., reported 2.1g of SDF and 2.7g of IDF per 100g in Hass avocados (Wang et al., 2015), which was higher than our results (1.59g and 2.09g, respectively, Table 2). Further, Li et al. analyzed the low-molecular-weight sugars, SDF and IDF of 70 foods, including avocados, which laid the foundation for many dietary fiber values reported in the USDA Food Database (B. W. Li et al., 2002). In that study, the researchers assessed California Hass avocados and Florida’s Fuerte cultivar. Interestingly, the total fiber (g/100g) was highest for Fuerte avocados (6.72g) compared to Hass avocados (5.53g). Consequently, the soluble (2.03g) and insoluble fiber (3.51g) were higher for Hass avocados compared to the results presented herein (1.59g and 2.09g, respectively, Table 2). The moisture level and the %DM differed for both varieties. Hass avocados had lower moisture (64.59% vs. 79.22%), and thus a higher %DM (35.41% vs. 20.78%), which translates to a higher fiber concentration expressed on a wet basis. The lower fiber content in Hass avocados compared to Fuerte indicates actual differences in fiber between varieties, rather than differences in moisture. However, within the same cultivar, differences in moisture can affect DF content. For instance, (Slater et al., 1975) compared dietary fiber from Hass avocados ripened in April and June, reporting that the avocados from June had higher DF than the ones harvested in April (2.80g vs. 2.20g/100g of fresh weight), possibly related to differences in moisture and %DM between months.

The analysis of DF across studies needs to consider the methods employed (e.g., enzymatic-gravimetric, enzymatic-chemical method, or enzymatic-gravimetric-chemical approach). The USDA primarily employs enzymatic-gravimetric methods, such as AOAC 985.29 and 991.43 (B. W. Li et al., 2002; Phillips et al., 2019; U.S. Department of Agriculture, 2014), which measure only SDF and IDF. In contrast, the Food and Drug Administration promotes the use of AOAC methods 2009.01 and 2011.25, which measure non-digestible oligosaccharides in addition to SDF and IDF (Food and Drug Administration, 2014, 2020). Thus, DF analyses of the same food may differ depending on the method used. Lastly, methodological differences in fat extraction (e.g., solvents such as hexane, diethyl ether, Folch) can influence fiber results, as samples with >10% fat, such as avocados, require fat extraction before analysis (Ford et al., 2023). In general, our TDF value (3.68g/100g) is more closely related to the pooled mean data from the literature (3.87g/100g) (Ford et al., 2023) compared to previous literature and food databases.

Despite the valuable insights gained, this study has several limitations. First, avocado’s food matrix—rich in MUFA, fiber, and micronutrients within a water-based emulsion—enhances fiber bioaccessibility but may interfere with fiber analysis (Dreher & Davenport, 2013). Second, the unique ripening behavior of avocados, compared to other climacteric fruits, and the variability in cultivars, postharvest handling, and seasonal factors complicate the DF analysis standardization (Brummell, 2006; Pedreschi et al., 2019). Third, the cell wall polysaccharide composition is challenging to resolve; the current method is not as sensitive to distinguish individual pectic polymers, as overlapping monosaccharide structures are found in pectin and hemicellulose. More refined techniques (e.g., esterification degree, molecular weight) may provide greater specificity than colorimetric uronic acid assays (Vierhuis, 2002). Lastly, the small sample size is a limitation. Future work should analyze multiple batches and avocado cultivars (e.g., Fuerte, Hass, Lula, Criollo) across ripening stages.

Nonetheless, this study has several strengths. It provides updated, detailed data on avocado DF components, distinguishing between SDF and IDF, and their respective monosaccharide profiles. This enabled the quantification of pectin, hemicelluloses, and cellulose fibers, which have known metabolic and gastrointestinal health benefits (An et al., 2022; Gill et al., 2021). The enzymatic-chemical-gravimetric method used is reproducible and comparisons can be made across avocado types and conditions. Importantly, our work is translationally relevant as results are reported on an“as is” basis.

## 1.4 CONCLUSIONS

This study aimed to characterize the amount and types of DF (e.g., pectin, hemicellulose, cellulose) and lignin in ripe avocados and evaluate the changes in DF at different stages of ripeness. The results of this study suggest ripe avocados contain a balanced amount of 43% SDF and 57% IDF. Of the SDF, RG-1 pectin is the main component with arabinan and arabinogalactan sidechains, whereas cellulose (32%), followed by hemicelluloses (23%) (e.g., xylans and xyloglucan), and lignin (2%) are the main components of IDF. As hypothesized, avocado ripening led to changes in DF, particularly a decrease in overall TDF, increased solubilization of pectin monosaccharides, and little to no changes in hemicellulose and cellulose. This study expands the database of DF in avocados, a nutrient-dense food that may help address the public health concern of inadequate fiber intake in the U.S. population.

## Supporting information

Article highlights

## CRediT authorship contribution statement

The author’s responsibilities were as follows:

**María G. Sanabria-Véaz:** Research design, Writing – Original draft, Methodology, Investigation, Formal analysis. **Knud Erik Bach Knudsen:** Methodology, Validation, Data curation, Writing – Review & Editing. **George C. Fahey:** Methodology, Validation, Writing – Review & Editing**. Hannah D. Holscher:** Research design, Resources, Writing – review & editing, Supervision, Project administration, Methodology, Funding acquisition.

## Declaration of Generative AI and AI-assisted technologies in the writing process

The authors used ChatGPT and Grammarly solely for editorial support to improve clarity, grammar, and readability of author-written text. After using this tool/service, the author(s) reviewed and edited the content as needed and take full responsibility for the content of the publication

## Acknowledgments

The following researchers are acknowledged for their contributions to this work:

Laura Bauer^2^ and Ryan Dilger^2^: Fat extraction methodology, “As-is” basis calculations, Review and editing of methods

## Abbreviations

%DM Abs: absolute dry matter percentage
DF: dietary fiber
DMB: dry matter basis
HG: homogalacturonan
IDF: insoluble dietary fiber
I-NCP: insoluble non-cellulosic polysaccharides
LMW: low-molecular weight
NSP: non-starch polysaccharide
RG-1: rhamnogalacturonan-1
RG-2: rhamnogalacturonan-2
SDF: soluble dietary fiber
S-NCP: soluble non-cellulosic polysaccharides

---

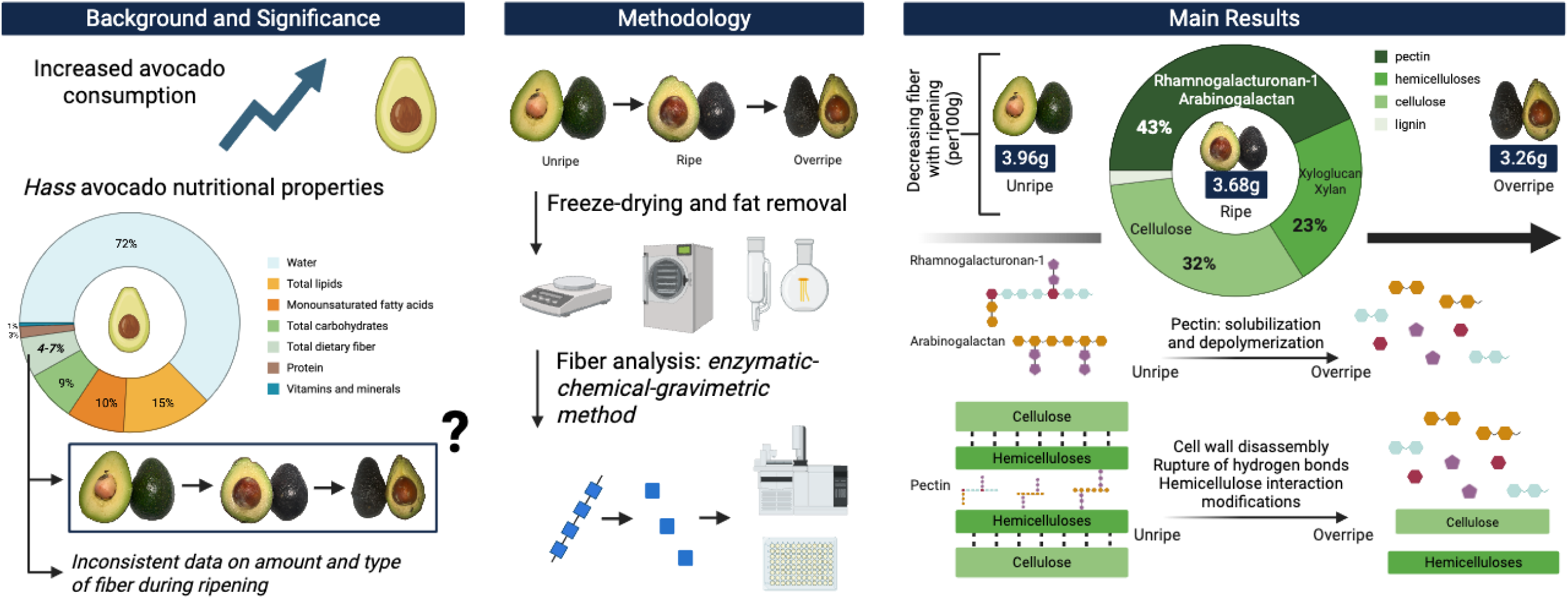

