## Supplementary files for "Quantifying and Characterizing the Fiber in Hass Avocados During the Ripening Process"

### Supplementary Materials

*Supplemental Table 1. Changes in avocado fiber constituents on a dry matter basis (DMB) at different ripening stages.*

| Avocado Ripening Stage | | NCP (%) | Cellulose (%) | NSP (%) | Lignin (%) | DF (%) |
|---|---|---|---|---|---|---|
| Unripe Day 0 | Soluble | 15.6 | | 15.6 | | 15.6 |
| | Insoluble | 14.3 | 13.3 | 27.9 | 1.0 | 28.6 |
| | Total | 30.0 | 13.3 | 43.3 | 1.0 | 44.3 |
| Ripe Day 5 | Soluble | 20.4 | | 20.4 | | 20.4 |
| | Insoluble | 10.8 | 15.1 | 25.9 | 0.9 | 26.8 |
| | Total | 31.2 | 15.1 | 46.3 | 0.9 | 47.1 |
| Overripe Day 12 | Soluble | 14.2 | | 14.2 | | 14.2 |
| | Insoluble | 9.9 | 16.6 | 26.5 | 1.0 | 27.5 |
| | Total | 24.1 | 16.6 | 40.7 | 1.0 | 41.7 |

*NCP = non-cellulosic polysaccharides, NSP = non-starch polysaccharides; DF = dietary fiber

*Supplemental Table 2. Non-cellulosic polysaccharide (NCP) residues on a Dry Matter Basis (DMB) from soluble and insoluble fractions of avocado fiber at different ripening stages.*

| Avocado Ripening Stage | | Rha (%) | Fuc (%) | Ara (%) | Xyl (%) | Man (%) | Gal (%) | Glu (%) | U.A. (%) |
|---|---|---|---|---|---|---|---|---|---|
| Unripe Day 0 | Soluble | 0.3 | 0.0 | 2.7 | 0.1 | 0.0 | 1.9 | 0.1 | 10.5 |
| | Insoluble | 0.2 | 0.4 | 2.6 | 3.6 | 1.3 | 2.5 | 1.1 | 2.8 |
| | Total | 0.5 | 0.4 | 5.3 | 3.6 | 1.4 | 4.4 | 1.2 | 13.3 |
| Ripe Day 5 | Soluble | 0.6 | 0.1 | 5.0 | 0.1 | 0.1 | 1.4 | 1.0 | 12.3 |
| | Insoluble | 0.1 | 0.4 | 1.7 | 4.1 | 1.3 | 1.4 | 0.0 | 1.9 |
| | Total | 0.7 | 0.4 | 6.7 | 4.0 | 1.4 | 2.8 | 1.0 | 14.2 |
| Overripe Day 12 | Soluble | 0.7 | 0.0 | 4.2 | 0.2 | 0.2 | 1.4 | 1.2 | 6.6 |
| | Insoluble | 0.1 | 0.4 | 1.1 | 4.0 | 1.4 | 1.4 | 0.0 | 1.5 |
| | Total | 0.8 | 0.4 | 5.3 | 4.1 | 1.6 | 2.8 | 1.2 | 8.1 |

*Rha=rhamnose, Fuc=fucose, Ara=arabinose, Xyl=xylose, Man=mannose, Gal=galactose, Glu=glucose, U.A.=uronic acid
