## Supplementary material for "Quantifying and Characterizing the Fiber in Hass Avocados During the Ripening Process": Article highlights

- Ripe avocados contain 43% soluble and 57% insoluble fiber.
- Soluble fiber is rich in pectin; insoluble fiber is mainly cellulose, with some hemicelluloses and lignin.
- Total dietary fiber tends to decrease during avocado ripening.
